# Modeling the Brain as an Information Source: An Information-Theoretic Framework for Decoding Cognitive States from fMRI

**DOI:** 10.64898/2026.04.21.719919

**Authors:** Gonul Gunal Degirmendereli, Abdulla Ahmadkhan, Fatos T. Yarman Vural

## Abstract

This study explores the hypothesis that the anatomical regions of the brain can be modeled as complementary and interconnected information sources. We propose a novel framework for analyzing the dynamics of these interacting information sources during cognitive tasks using information-theoretic measures. Specifically, we introduce dynamic and static entropy models to quantify the information content within individual anatomical regions, both over time and in relation to specific cognitive demands. Furthermore, we develop two network models based on the dynamic and static Kullback-Leibler (KL) divergence to characterize the regional interactions.

Testing our models on fMRI data recorded during Complex Problem Solving (CPS) tasks reveals promising results. Entropy values successfully identify activated brain regions, consistent with the existing neuroscience literature. Furthermore, our Kullback-Leibler network models demonstrate high accuracy in distinguishing between the planning and execution phases of CPS, as well as in differentiating between expert and novice problem solvers. These findings suggest that our information-theoretic approach holds promise for identifying active brain regions, characterizing mental states, and elucidating brain networks associated with cognitive tasks.

## 1 Introduction

Recent advances in brain research suggest that the brain’s anatomical regions can be conceptualized as parallel and dynamic information-processing units that interact to form complex networks. This perspective allows us to use information theory (Shannon, 1948) as a framework to quantify and characterize the information flow across brain anatomical regions to decode cognitive states. Although cognition is too complex to be explained by a single framework, information theory offers powerful tools for analyzing and interpreting fMRI signals recorded during specific cognitive tasks. Specifically, the concepts of *self-information* and *entropy* are very suitable to quantify the information generated in anatomic regions.

Generally speaking, *self-information* measures the information content of a random event, reflecting the uncertainty about a single outcome. On the other hand, *entropy* measures the average amount of information for all possible outcomes generated by a random source. It reflects the randomness within a system and indicates the level of uncertainty associated with those outcomes. In other words, entropy represents the degree of incompleteness in our knowledge about the precise state of the system under consideration (Bavaud et al., 2005; Cengel, 2021).

A fundamental insight into neural computation is that uncertainty is an inherent feature of brain information processing. Whether in processing sensory input, controlling movement, or abstract reasoning, neural computation inherently deals with uncertainty (Pouget et al., 2013). Identifying how the brain represents and operates under uncertainty is key to understanding how it makes inferences. An effective strategy for reasoning under uncertainty is to represent information using probability distributions and to make predictions based on probabilistic inference rules (Van Horn, 2003; Fiser et al., 2010; Yuille & Kersten, 2006; Pouget et al., 2013).

Information theory thus offers a natural and efficient method for explaining how neural systems use probability calculations to manage uncertainty. There is substantial support from neuroscience for using entropy to measure neural activity, offering insights into the probabilistic nature of brain function (McIntosh et al., 2008; Lizier et al., 2011; Wang, Z. et al., 2014; Viol et al., 2017; Gunal Degirmendereli, & Yarman Vural, 2021). Consequently, entropy can be utilized to decode mental states by identifying characteristic patterns of brain activity associated with different mental states.

This study introduces an information-theoretic model to represent a fundamental cognitive ability: Complex Problem Solving (CPS). CPS is not only seen as a valuable skill for individuals but is also seen as vital to the future of societies (OECD, 2023; World Economic Forum, 2023; McKinsey, 2023). As the world becomes increasingly interconnected and technologically advanced, the convergence of digital, physical, and biological systems produces complex challenges that require advanced problem-solving skills (Schwab, 2017).

Recent studies in information-theoretic neural signal analysis highlight the importance of dynamic, time-resolved connectivity metrics for capturing nonstationary brain states. Studies such as Faes et al. (2016) and Antonacci et al. (2025) have shown that entropy and divergence measures can reveal meaningful temporal changes in neural activity, particularly in EEG data. Faes et al. (2016) employed time-varying entropy and mutual information to quantify dynamic neural interactions, while Antonacci et al. (2025) demonstrated that local, time-resolved information-theoretic metrics can effectively capture transient neural dynamics during cognitive tasks. Extending this line of work, Antonacci et al. (2025) applied time-varying information-theoretic measures to model nonstationary neural processes more broadly, reinforcing the value of dynamic entropy frameworks.

Building on these principles, we introduce a novel application of time-resolved Kullback–Leibler (KL) Divergence to fMRI data. By treating each anatomical region as a dynamic information source, we track evolving probability distributions over voxel intensities and quantify inter-regional divergence across time. This framework enables a spatially grounded, temporally resolved analysis of functional reorganization during cognitively demanding tasks—expanding the scope of information-theoretic analysis into the domain of high-resolution fMRI.

Our computational model is based on the premise that the fMRI signals recorded during CPS tasks originate from a multitude of random sources within the brain. These sources, corresponding to distinct anatomical regions, can be effectively modeled using the mathematical framework of Shannon information theory. fMRI signals reflect brain activity patterns that represent the mental states induced by the task, encapsulating information about their expected probabilities of occurrence.

In the proposed model, mental states are represented by the amount of information generated within anatomical regions and the communication of information between them. The entropy value characterizes the amount of information within the anatomical regions, while the KL divergence quantifies the flow of information between them (Gunal Degirmendereli & Yarman Vural, 2022).

First, we estimate the probability density functions of the fMRI signals from each anatomical region to quantify their information content during CPS. From these distributions, we define static and dynamic entropy measures, capturing the average information content of each source in different problem-solving phases.

Furthermore, we construct static and dynamic brain networks to explore interregional interactions during CPS, comparing skilled and less skilled problem solvers. We validate our computational models by comparing them with findings from experimental neuroscience. Finally, we conduct a series of brain decoding experiments to decode the phases of CPS. We compare the decoding performance of our models with those based on Pearson correlation and raw fMRI BOLD signals. Notably, our entropy- and KL-divergence-based network models outperform conventional multi-voxel pattern analysis (MVPA) models (Alchihabi et al., 2021).

## 2 Modeling the Human Brain as a Set of Information Generation Sources

In this section, we model the brain as an information generation and communication system using Shannon information theory, which quantifies the average information exchanged through neuronal spikes (Oweiss, 2010), assuming stochastic and noisy coding and transmission. The Shannon–Weaver communication model (Weaver, 1953) includes five components:

1. An **information source** generates messages.
2. A **transmitter** converts the message into a transmittable signal.
3. The **channel** transmits the signal, which is potentially affected by noise.
4. The **receiver** decodes the signal to retrieve the message.
5. The **destination** is the intended recipient.

Neuronal communication, the essential process underlying brain function, can be conceptualized using Shannon’s framework of communication over a noisy channel, which helps to clarify the challenges and mechanisms involved in neural processes. These approaches allow us to draw the following parallels with information processes in the brain:

1. **Information source and Receiver**: Each anatomical region can be considered as a Shannon information source that generates and transmits information to related regions. In this context, an anatomical region is viewed as both a source and a receiver.
2. **Message Encoding, Transmitting and Decoding**: In neural processing, information is encoded as neural activity, transmitted via synapses, and decoded by downstream neurons, akin to Shannon’s encoding, transmission, and decoding.
3. **Noise and Error Correction in the Channel**: Just as Shannon’s model incorporates noise and error correction, neural signal transmission is influenced by interference, with synaptic plasticity, playing a key role in error correction and adaptive adjustment within the communication channel.
4. **Bandwidth and Capacity**: Similar to Shannon’s model for channel bandwidth and capacity, neural networks operate within constraints on information transmission bandwidth and synaptic processing capacity.

Although there are significant differences between communication systems and neural systems, these analogies facilitate insights into the fundamental principles governing information processing in both areas.

An fMRI signal provides noisy information about neuronal activation within a collection of neurons, referred to as a voxel (a cubic brain volume). Experimental neuroscience indicates that collections of voxels within anatomical regions exhibit coordinated activity, during cognitive tasks. We hypothesize that a mental state is generated from highly correlated group of anatomical regions, each of which is modeled as an information source. fMRI signals reflect the brain activity patterns and convey information about mental states.

Measuring the information content of brain anatomical regions enables us to infer their states as outcomes of cognitive tasks. Information theory suggests that signals from an information source have low entropy when a specific outcome is highly probable, and high entropy when all potential outcomes are equally likely.

Figure 1 illustrates the proposed computational model for decoding mental states. In this model, each information source (anatomical region) generates a sequence of messages embedded in noisy fMRI signals. The information content of the fMRI signals from each information source is measured by Shannon entropy to predict a mental state as the outcome of a cognitive task. We expect that anatomical regions with structured patterns produce low entropy values, indicating a specific mental state. In contrast, regions with random fluctuations would exhibit high entropy values, indicating nonparticipation in the task. (Gunal Degirmendereli et al., 2019).

**Figure 1:**
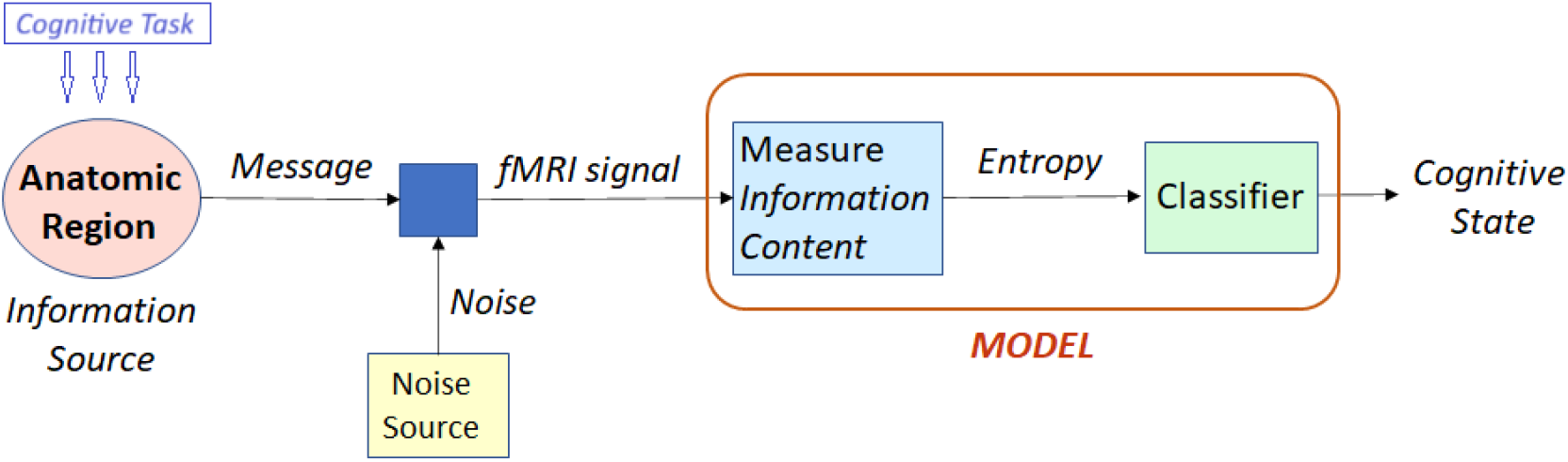
The proposed computational model for decoding mental states by treating anatomical regions as information sources.

Subsequently, we propose a method to estimate static and dynamic brain networks among anatomical regions using relative entropy, known as Kullback-Leibler (KL) divergence. We assume that the degree of co-activation between two anatomic regions can be measured by KL divergence. We also assume that highly co-activated regions have relatively lower KL divergences compared to non-activated regions (Gunal Degirmendereli, & Yarman Vural, 2021). This allows us to define brain networks, where nodes represent anatomical regions, and arc weights denote the KL divergence values between them. Figure 2 illustrates the proposed computational model designed to estimate the connectivity between the anatomical regions and to decode brain states associated with a cognitive task.

**Figure 2:**
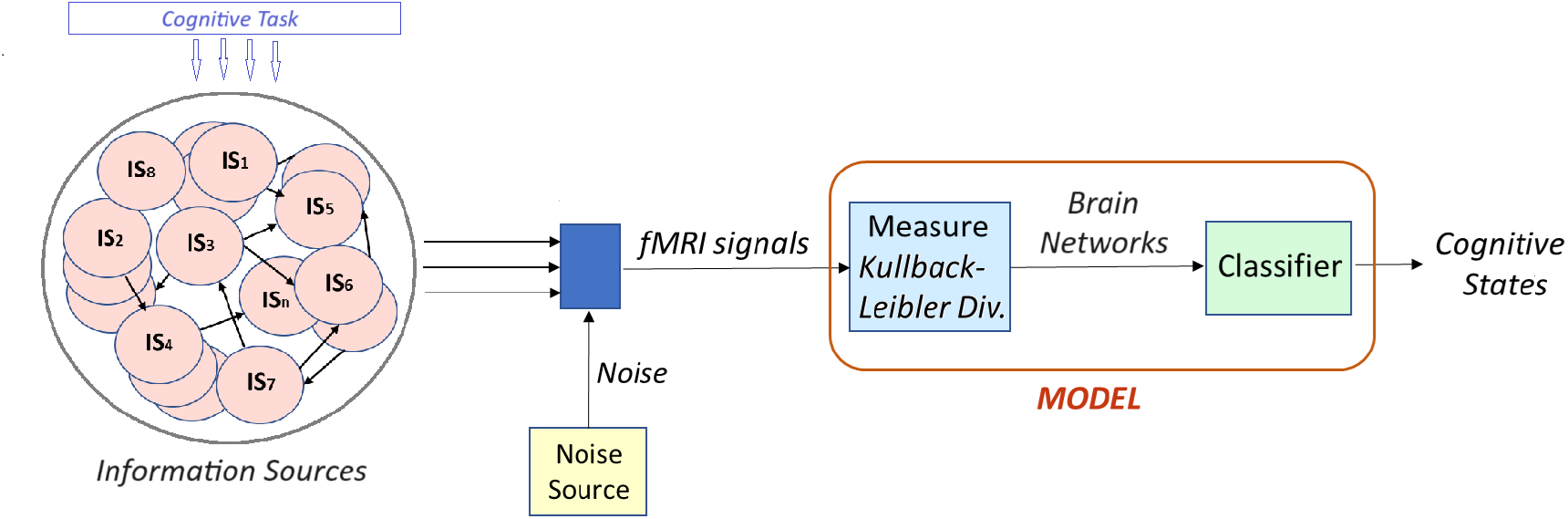
Proposed computational model for estimating the connectivity among the anatomical regions, modeled as information sources (IS). The connectivity between the anatomic regions are estimated by KL divergences, which are used as feature vectors for decoding brain states associated with a cognitive task.

### 2.1 Measuring the Information

Let us formally define the major concepts, mentioned above, namely self-information, information entropy, and Kullback-Leibler divergence, as follows:

#### Self-Information

Consider a probabilistic event, represented by a discrete random variable *x*_*i*_, with all possible outcomes 𝒳 = {*x*_1_, *x*_2_, …, *x*_*n*_}, and the associated probability mass function, *P* (*x*_*i*_). Self-information for the outcome *x*_*i*_ is defined as:

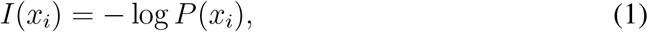

where log is the logarithm, and *P* (*x*_*i*_) is the probability of the outcome *x*_*i*_, out of all possible outcomes of the event. Intuitively, −log *P* (*x*_*i*_) is proportional to the uncertainty of the outcome, *x*_*i*_. When the base of the logarithm is 2, the unit of self-information is bits (binary units).

#### Entropy

The average amount of information conveyed by an event, when considering all possible outcomes within a system, is measured by entropy. It is defined as the expected value of self-information.

Formally, for a discrete random variable *x*_*i*_, the entropy is defined by taking the expected value of Eq. 1 over the set of all possible outcomes, 𝒳:

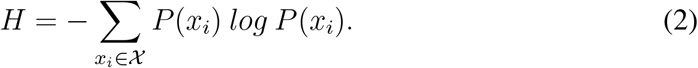

The unit of entropy is bits per anatomical region for the base-2 logarithm. It is zero only when all but one *P* (*x*_*i*_) are zero, indicating certainty. Otherwise, *H* is positive. *H* reaches its maximum when all *P* (*x*_*i*_) are equal, representing maximum uncertainty. In this study, we use the base-2 logarithm for all definitions.

#### Kullback-Leibler Divergence

Consider two probability density functions, *P*_*k*_(*x*_*i*_) and *P*_*l*_(*x*_*i*_), defined over a random variable, *x*_*i*_. Kullback–Leibler divergence between these two probability density functions is defined as,

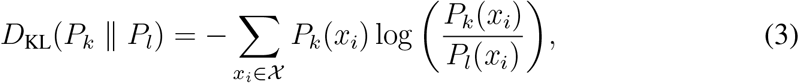

where 𝒳 is the set of all possible outcomes. Equation 3 shows the expectation of the difference between the self-information of the random event represented by *P*_*k*_(*x*_*i*_) and *P*_*l*_(*x*_*i*_), where the expectation is taken over the probability distribution function, *P*_*k*_(*x*_*i*_).

## 3 Data Generation Method: Experimental Setup and Nature of fMRI data

In this study, we focus on the Tower of London (TOL), a widely recognized game used to assess executive functioning and planning ability in complex problem-solving tasks. The fMRI data were recorded while 18 subjects (aged 19–38) played a computerized version of the TOL (Newman et al., 2009).

Each puzzle consisted of two configurations: initial and goal states. Participants were instructed to transform the initial state into the goal state using the fewest possible moves. Problem complexity was determined by the number of required moves, which varied based on the initial and goal positions. In this experiment, each puzzle required a minimum of five or six moves to reach the goal configuration.

The cognitive experiment consists of two phases:

1. **Planning Phase**: The subjects generated a solution plan before making their first move. This phase lasted from the onset of the puzzle presentation until the first move.
2. **Execution Phase**: The subjects executed their plan by moving a cursor to solve the puzzle. This phase began with the first move and continued until the puzzle was completed.

Each subject underwent four scanning sessions, each containing 18 puzzles with a 15-second time limit per puzzle. The initial 5 seconds were designated for planning, although subjects could extend this phase if they had not finalized their solution plan. Subjects were categorized based on their puzzle-solving performance. Those who successfully solve the most puzzles are labeled as *expert players* (12 subjects), while the remaining subjects are termed *novice players* (6 subjects).

The fMRI data comprised 590 time instants per session, yielding a total of 72 sessions and 1,296 puzzles across all subjects. The recorded data included 116 anatomical regions of interest defined by the Anatomical Automatic Labeling (AAL) atlas from the Montreal Neurological Institute (Tzourio-Mazoyer et al., 2002). However, the Cerebellum and Vermis regions were excluded, and the rest 90 regions were analyzed.

The fMRI images were acquired using a 3T Siemens TRIO scanner with an 8-channel radiofrequency coil at the Imaging Research Facility, Indiana University. The imaging parameters were: TR = 1000 ms, TE = 25 ms, flip angle = 60, voxel size = 3.125 × 3.125 × 5 mm with a 1 mm gap, across 18 slices of 5 mm thickness.

The fMRI dataset underwent a standardized preprocessing pipeline using statistical parametric mapping (SPM2) including slice acquisition timing correction (resampled to 2×2×2 mm voxels), spatial normalization (Gaussian filter of 8 mm), high-pass filtering (1/128 Hz cutoff frequency to remove low-frequency signals), and motion correction” (Newman et al., 2009).

The fMRI recordings were structured as four-dimensional space-time data, where intensity values represented neural activity at each voxel coordinate, *v*_*i*_(*t*) = (*x*_*i*_, *y*_*i*_, *z*_*i*_) are measured at each time instant, *t*. As a result, the data consisted of time-series, *v*_*i*_(*t*), recorded at each (*x*_*i*_, *y*_*i*_, *z*_*i*_) coordinate of a brain volume. In other words, at each time instant, the fMRI method captured a brain volume of intensity values, each associated with a corresponding cognitive state label:*planning or execution* (Gunal Degirmendereli, & Yarman Vural, 2021).

## 4 Estimating Dynamic and Static Entropy

In this section, we investigate the information content of brain anatomical regions during the *planning* and *execution* phases of complex problem solving, distinguishing between expert and novice players, using first-order Shannon entropy as a metric. Our major assumption is that the regions with lower entropy levels are more intimately involved in complex problem solving processes than regions exhibiting higher entropy. Therefore, we investigate the relationship between the phases of problem solving tasks and the entropy measures of anatomical regions. The voxel intensity values, measured under a cognitive state are regarded as random variables, recorded through fMRI scans as subjects solve complex TOL problems.

We introduce two entropy estimation methods, static and dynamic entropy, to quantify the information content present in signals originating from brain regions, as described below.

- **Dynamic entropy** is defined as the variation of the information content of the anatomical regions as a function of time. We define the intensity value of voxel *i*, at time *t* as a random variable, *v*_*i*_(*t*).
- **Static entropy** is defined over a period of time while the subject is resting or solving a TOL problem. We represent each anatomic region by the average BOLD signal of voxels in that region. Then, we define each planning or execution time instant of the average BOLD signal as a random variable.

### 4.1 Dynamic Entropy Estimation for Anatomic Regions

In this approach, we assume that the entropy of an anatomical region varies as a function of time. These variations can provide information about the CPS process and its planning and execution phases.

In order to estimate the dynamic entropy of an anatomical region, we use voxel intensity values, *v*_*i*_(*t*), measured at location (*x*_*i*_, *y*_*i*_, *z*_*i*_) at time *t*. For each time instant, the probability distribution of a region is estimated, assuming that the voxel intensity values within that region are random variables.

Let, *s* shows the index of an information source, which represents an anatomical region, where *s* ∈ {1, 2, 3, …, *S*} and *S* is the total number of regions; *v*_*i*_(*t*) is the voxel intensity value measured at coordinate (*x*_*i*_, *y*_*i*_, *z*_*i*_) in the anatomical region, *s*, at time *t*, where *i* ∈ {1, 2, …, *n*_*s*_}, and *n*_*s*_ is the number of voxels in the region *s*.

For each time instant, recorded in an fMRI session, the probability density function of the voxels in an anatomical region, *P*_*s*_(*v*(*t*)) is estimated using the kernel probability density estimation method, from the following equation:

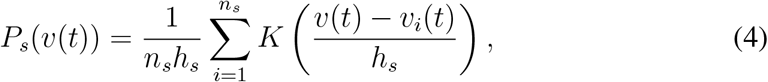

where *K* is the Gaussian kernel function and *h*_*s*_ is the smoothing bandwidth of the kernel function for source *s*. The standard deviation, *h*_*s*_, for the anatomic region, *s*, is called the *bandwidth*, which is a crucial parameter to be selected. The bandwidth has a strong influence on the estimated probability density function. Instead of setting a single global bandwidth, we estimate different bandwidth values for each source, based on the characteristics of voxel intensity values.

Using the aforementioned approach, we represent each anatomical region as an information source, characterized by two-tuples, [*v*(*t*) ∈ **s**, *P*_*s*_(*v*(*t*)]. This representation allows us to estimate a dynamic entropy measure for each source, **s** as a function of time, *t*, as follows;

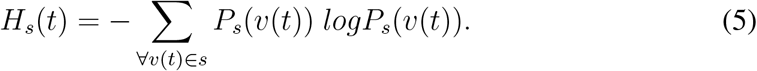

### 4.2 Static Entropy Estimation for Planning and Execution Phases at Each Anatomic Region

In this section, we estimate a scalar entropy value for each anatomical region during the planning and execution phases, separately. The static entropy of an anatomical region is considered as a measure of its contribution to either the planning or execution phase. This measure allows us to compare our proposed model with findings from experimental neuroscience, which identifies active anatomical regions for each phase. In this approach, we assume that each anatomical region can be represented by a time series derived from the expected value of all voxel time series within that region. Then, we define the static entropy of the information source, based on the random variable, defined by each instant of the representative time series.

Consider an fMRI time series, recorded during a CPS session, *v*_*i*_(*t*) at each voxel coordinate *i* to represent the neural activity of the underlying CPS task. The representative time series, *X*_*s*_(*t*) for an anatomical region, **s**, is estimated by taking the expected value of all of the time series, which resides in that region,

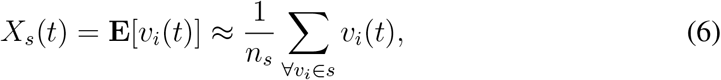

where **E** is the expectation operator, *n*_*s*_ is the number of voxels in the anatomical region, represented by the source **s**.

To characterize the information content of the planning and execution phases, for each source *s*, we estimate two probability density functions: One for the planning phase and the other one for the execution phase, using the label of each time instant of fMRI data. The probability density functions in source **s**, for *planning* phase *P*_*ps*_(*x*) and *execution* phase *P*_*es*_(*x*), are estimated using the kernel density estimation method, as follows:

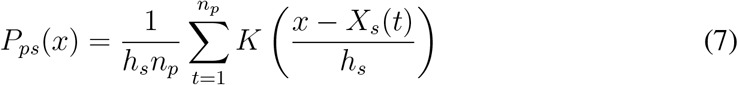

and

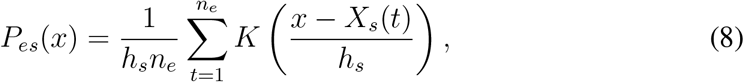

where *n*_*p*_ and *n*_*e*_ are the number of time instants for planning and execution phases respectively. *X*_*s*_(*t*) is the representative time series for a source *s. K* is the Gaussian kernel smoothing function, and *h*_*s*_ is the bandwidth value, determined by the method, suggested in (Silverman, 1986).

Once we estimate the probability distribution functions for the source **s**, for *planning* phase *P*_*pl,s*_(*x*) and *execution* phase *P*_*ex,s*_(*x*), the static entropy of the information source **s** for *planning* phase is then estimated by,

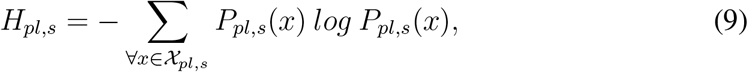

where 𝒳_*ps*_ is the set of samples in anatomical region *s*, which belong to the planning time instants.

Similarly, the static entropy of the information source **s** for *execution* phase is then estimated by,

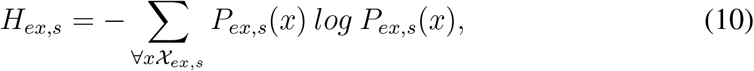

where 𝒳_*ex,s*_ is the set of samples in anatomical region *s*, which belong to the execution time instants.

Note that, in the above formulations, representative fMRI time series *X*_*s*_(*t*), of each information source, for all sessions of a subject were concatenated, separately for *planning and execution* phases. Then, the probability distributions were calculated for *planning and execution* phases for each subject and each information source.

### 4.3 Static Entropy Estimation for Expert and Novice Players

In this section, we estimate a scalar entropy value for each anatomical region for *expert* and *novice* players during the planning and execution phases. This metric allows us to compare the level of randomness in anatomical regions between expert and novice players, when performing planning and execution tasks

We estimate the static entropy separately for planning and execution phase samples using the voxel time series from the underlying source. Our approach involves initially estimating dynamic entropy for each anatomical region and for each time instant of planning and execution, using the distribution of voxel intensity values within that region. Subsequently, we segregate the successful and unsuccessful runs. Finally, we derive scalar entropy values for each region by averaging across the time instants of expert players and novice players, respectively, as they engage in their planning and execution tasks.

Consider an anatomical region, represented by the information source, **s** for *s* = 1, …, *S*. We group all of time samples, *t* of the dynamic entropy function *H*_*s*_(*t*), defined in Equation 5 into four sets:

- *T*_*pl,su*_: The set of time samples, that belong to successful (*su*) runs, while the subject performs the planning (*pl*) task.
- *T*_*ex,su*_: The set of time samples, that belong to successful runs while the subject performs execution (*ex*) task.
- *T*_*pl,un*_: The set of time samples, which belong to unsuccessful (*un*) runs, while the subject performs the planning task.
- *T*_*ex,un*_: The set of time samples, that belong to unsuccessful runs while the subject performs the execution task.

Then, we estimate four scalar entropy values for each anatomical region, **s**, from the following equation:

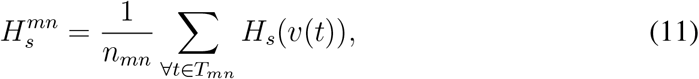

where *m* ∈ {*pl, ex*}, *n* =∈ {*su, un*} and *n*_*mn*_ is the number of time instants in set, *T*_*mn*_.

## 5 Estimating Static and Dynamic Kulback-Leibler Divergences

Representation of anatomical regions as a set of information sources allows us to estimate the connectivity among these sources by Kullback-Leibler (KL) divergence. KL divergence quantifies the statistical dissimilarity between the probability distributions of fMRI signals from different brain regions, enabling the construction of network representations that capture evolving neural interactions during cognitive tasks.

Our key assumption is that the degree of co-activation between two brain regions, modeled as information sources, can be assessed using KL divergence. The KL divergence of 0 indicates that the two distributions are identical. The KL divergence between pairs of anatomical regions serves as the arc weights in the brain network, defining the connectivity among these information sources.

Based upon this assumption, we propose two types of KL divergences, depending on the random variable defined for each information source:

- **Static KL divergence** between two anatomical regions is estimated from the probability distribution function of each region by taking the expected value over the time instants of planning and execution phases, separately. It is used to form static brain networks to investigate the co-activation of anatomical regions for the planning and execution tasks.
- **Dynamic KL divergence** between two anatomical regions is estimated from the probability distribution of voxel time-series, at each time instant. It is used to form dynamic brain networks to investigate the co-activation of regions as a function of time.

### 5.1 Static Kullback-Leibler Divergence Estimation

In order to estimate static brain networks, we define each time instant of the representative time series, *X*_*s*_(*t*), at each anatomic region, as a random variable. Then, we estimate a probability distribution function for each anatomic region and for each cognitive state, namely, *planning and execution*.

Let us formally describe the method for estimating static Kullback-Leibler divergence to represent the *planning* and *execution* phases for each anatomical region. Recall that, the representative time series, *X*_*s*_(*t*) for an information source **s** is estimated by (6), described in Section 4.2.

For each anatomic region represented by the source *s*, we estimate two probability distribution functions: One for the *planning* phase and the other one for the *execution* phase, using the label of the time instants. The probability distribution functions in source *s*, for *planning* phase *P*_*pl,s*_(*x*) and *execution* phase *P*_*ex,s*_(*x*), are estimated by Equations (7) and (8) described in Section 4.2.

When we consider the time instants of a representative time series with *planning* label, the corresponding static KL divergence between the probability distributions of two anatomical regions, modeled by the information sources *k* and *l*, for ∀ *l* ≠ *k* is, then, estimated by

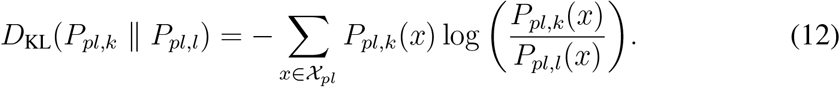

where 𝒳_*pl*_ is the set of all samples, which belong to the planning phase. On the other hand, when we consider only the time instants with execution label, the static KL divergence is estimated by

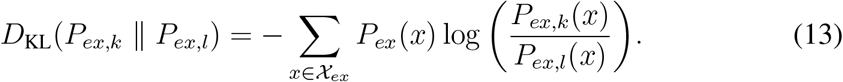

where 𝒳_*ex*_ is the set of all samples, which belong to the execution phase.

In the above formulations, the representative time series *X*_*s*_(*t*) for each anatomical region are concatenated across all sessions, separately for the *planning* and *execution* phases. The probability distributions are then estimated for both phases for each anatomical region.

### 5.2 Dynamic Kullback-Leibler Divergence Estimation

In order to estimate a dynamic brain network, we directly use voxel intensity values, *v*_*i*_(*t*), measured at source location (*x*_*i*_, *y*_*i*_, *z*_*i*_) at time instant, *t*. For each information source and time instant, the probability distribution function over all voxels in a source *P*_*s*_(*v*(*t*)) is estimated using (4), described in section 4.1.

The dynamic KL divergence between two anatomical regions, represented by the information sources *k* and *l* for ∀ *k* ≠ *l* at time *t* is estimated as follows;

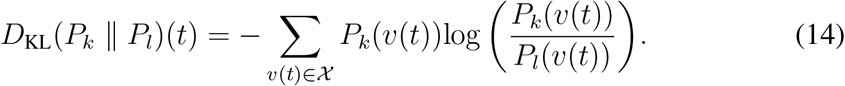

where 𝒳 is the set of voxel intensity values at a particular time instant *t*, in the anatomical region, *k* or *l*.

## 6 Dynamic and Static Brain Networks

Representing fMRI signals through brain networks is essential for comprehending the exchange of information among anatomical regions during cognitive tasks. The complexity of human behavior arises from the organization of neurons into anatomical circuits. Each neuron acts as a node within one or more networks and transmits information relevant to one or more specific cognitive tasks. It is widely recognized that neurons with similar characteristics contribute to the same brain activities depending on how they are connected with each other. The specific patterns of neural connections and how these circuits are functionally organized in the brain drive our thoughts and behaviors.

In this study, we define two types of brain networks across the anatomical regions using the static and dynamic KL divergences, introduced in the previous section. Static brain networks are defined for planning and execution phases, separately, as two tuples,

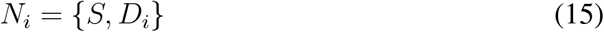

where each node, *s* ∈ *S* represents an anatomical region, and the edge weights between the anatomic regions *k* and *l*, 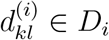 are the static KL divergences,

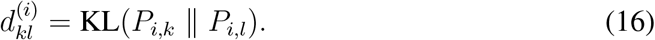

for planning and execution phases, *i* = *pl, ex*, respectively.

Similarly, the dynamic brain networks are defined as two tuples,

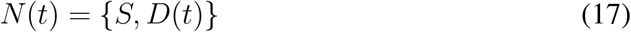

where the nodes *s* ∈ *S* are the same as the static network, and the edge weights *d*_*kl*_(*t*) ∈ *D*(*t*) are the dynamic KL divergences, between the anatomic regions *l* and *k*,

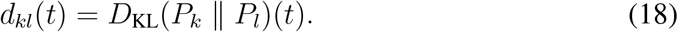

Note that, in the above formulation, the static brain networks are defined for planning and execution phases separately. Note also that the dynamic brain networks vary as a function of time.

## 7 Experiments on Static and Dynamic Entropy Representations

In this section, we examine the behavior of dynamic and static entropy variations of anatomical regions for expert and novice players, during the planning and execution phases of CPS.

### 7.1 Dynamic Entropy Variations of Complex Problem Solving and Resting States

In this part, we investigate the dynamic entropy variations within anatomical regions during the resting state and CPS task, identifying regions with low and high entropy. As a reminder, low entropy indicates organized and predictable behavior of the relevant source, while high entropy sources exhibit relatively random behaviors.

Initially, we estimate the dynamic entropy for each subject and each session, over the voxels that reside in an anatomical region. During the experiments, we notice that some subjects have relatively more random fluctuations of dynamic entropy, compared to some other subjects. We assume that the fluctuations of entropy, which deviate from the expected response to event-related stimuli, may serve as an indicator of an individual’s level of expertise.

Given the tendency for dynamic entropy measures among novice players to exhibit random fluctuations, our analysis is confined to the 12 expert players. Subsequently, we aggregate all dynamic entropy measures across 42 sessions involving these 12 subjects for each anatomical region and plot them over time.

For expert players, the measures of dynamic entropy are quite different in high and low entropy anatomical regions. Specifically, dynamic entropy tends to exhibit more structured fluctuations over time in low-entropy regions, while it shows random fluctuations in high-entropy regions. This fact is illustrated in Figure 3 and Figure 4. When analyzing these figures, it is important to account for a slight shift in the entropy plot (blue) compared to the experimental setup plot (red) due to the delay in the hemodynamic response to stimuli. Therefore, adjusting the experimental setup plot slightly to the right would align it more accurately with the entropy curve. This delay should be considered when examining entropy variations in subsequent sections.

**Figure 3:**
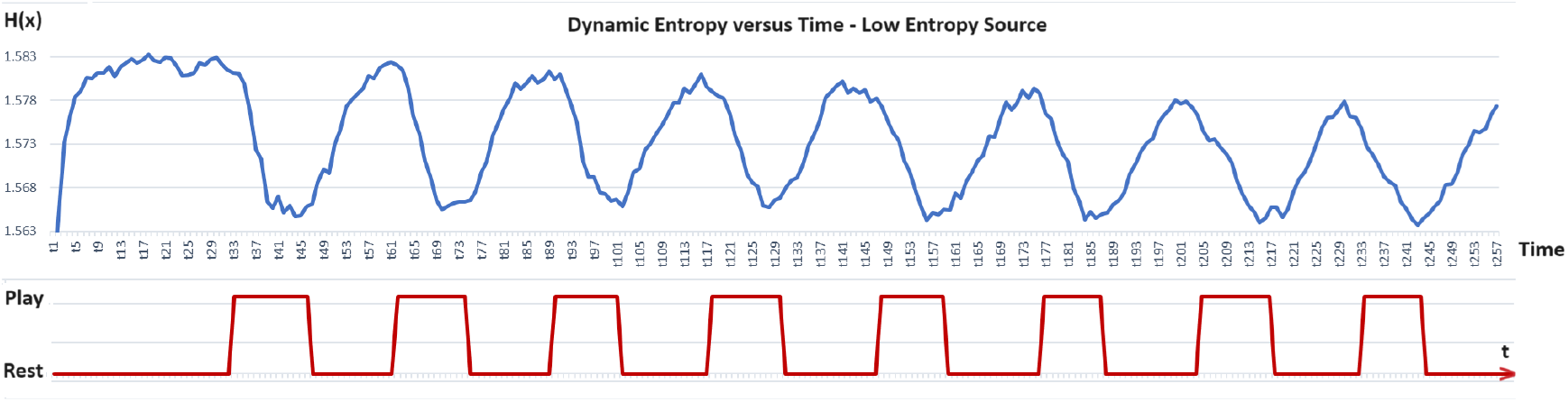
Dynamic Entropy vs. time for a minimum entropy source: Entropy fluctuations (blue) for *Left Superior Parietal* during TOL puzzle solving (averaged for 12 subjects, 42 sessions). *Play* shows the time instants when the subject plays the TOL game and *Rs* shows, when the subject is back to the resting state, in red plot. The first puzzle begins at 33. time-instant.

**Figure 4:**
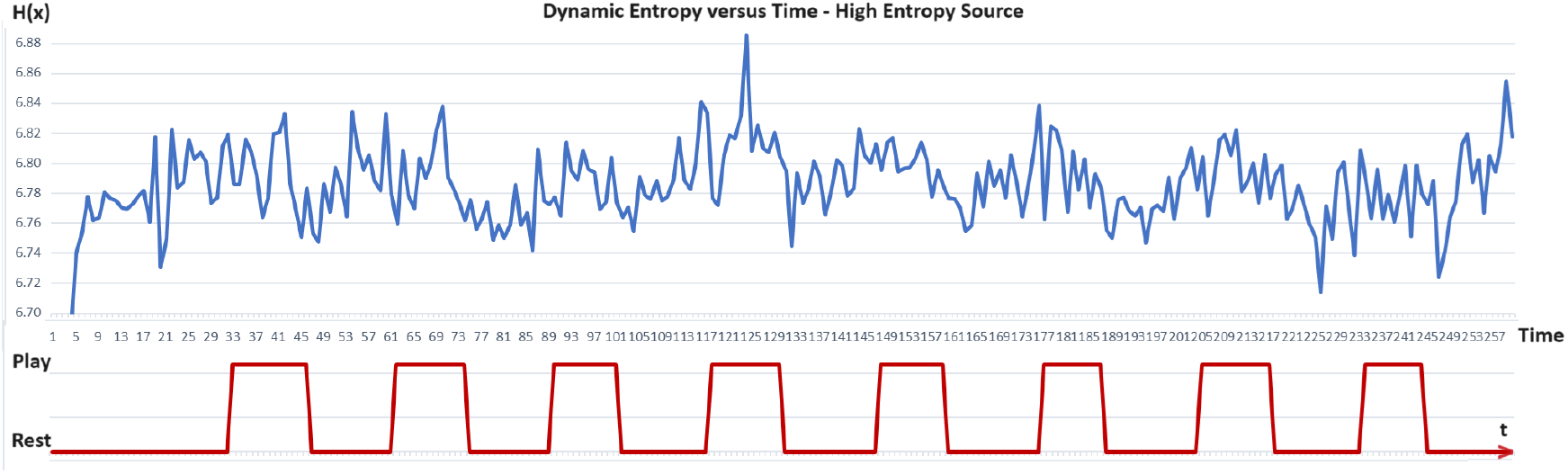
Dynamic Entropy vs. time for a maximum entropy region: Entropy fluctuations (blue) for *Right Insula* during a TOL session (averaged for 12 subjects, 42 sessions). *Play* shows the time instants, when the subject plays the TOL game and *Rs* shows, when the subject is back to the resting state, in the red plot.

As depicted in Figure 3, the dynamic entropy of regions with low entropy displays a strong correlation with the experimental setup. It consistently remains low during the TOL game and consistently high during rest periods. This suggests that playing the TOL game results in a more organized signal in low entropy regions. In simpler terms, a source of information that yields a more ordered outcome exhibits lower entropy values. “Order” here refers to a reduction in the number of states within the system, increasing certainty for a specific state. Thus, lower entropy values imply greater predictability of the source system’s state.

However, when examining dynamic entropy variations in high entropy regions, we observe irregular fluctuations, that appear to be heavily influenced by noise. As illustrated in Figure 4, dynamic entropy measures show little correlation with the experimental setup. As mentioned earlier, higher entropy values indicate greater unpredictability in the behavior or outcome of the system.

### 7.2 Dynamic Entropy Variations of Resting State, Planning and Execution Subtasks for Expert and Novice Players

In the first set of experiments, we investigate the dynamic entropy changes during the resting state, as well as the planning and execution phases of complex problem solving, for both expert and novice players. For this purpose, we partitioned the players into two groups based on the number of puzzles they successfully completed in a session and how many extra moves they made compared to the shortest path to complete a puzzle. We considered 12 subjects, who successfully completed 756 puzzles in a total of 42 successful sessions as *expert players* and the rest as *novice players*.

Then, we estimate the dynamic entropy measures of each anatomical region of each subject. Finally, we investigate the behavior of the dynamic entropy in the most informative (low entropy) anatomical regions for the expert and novice players.

For an *expert player*, the dynamic entropy during the resting state, and planning-execution phases is closely aligned with the experimental setup. (Figure 5). Specifically, high entropy values are observed during the resting state, which then drop sharply when the experts begin to play. This pattern aligns with the understanding that high entropy corresponds to a system occupying many states in its state space with equal probability, indicating randomness. Conversely, lower entropy values suggest more organized and structured activities. This same pattern is observed across all expert players. Therefore, dynamic entropy values indicate that expert players during play exhibit low-entropy brain regions corresponding to more structured and organized information generation compared to the resting state.

**Figure 5:**
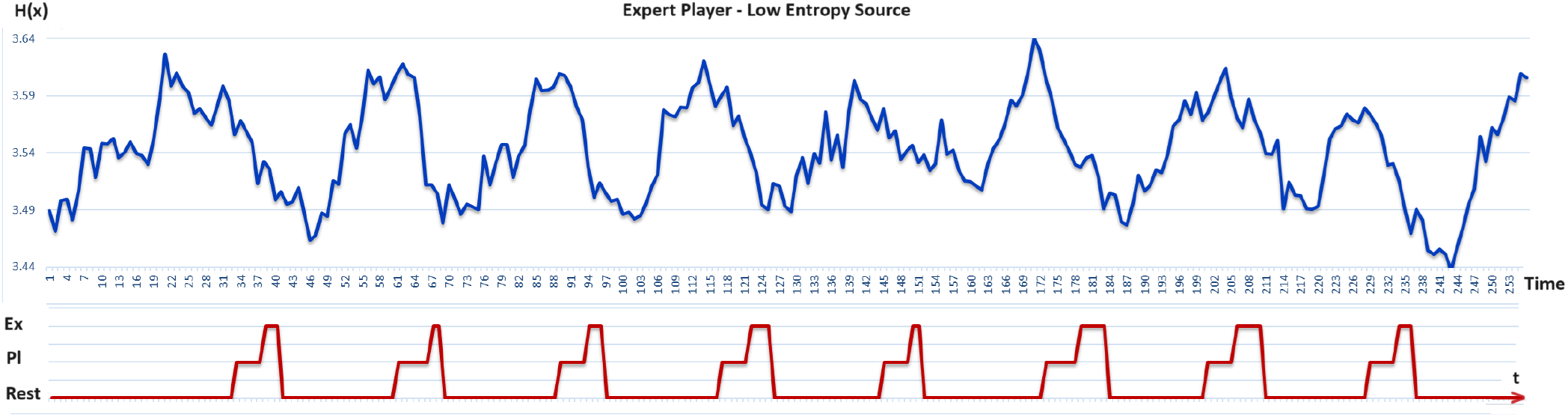
Expert Player: Dynamic Entropy fluctuations for a low entropy region, *Right Precuneus* (blue). *Pl* shows the Planning, *Ex* shows the execution and *Rs* shows the resting state time instants (red). (Gunal Degirmendereli et al., 2019)

For a *novice player*, the relationship between dynamic entropy and task performance is more random compared to expert players (Figure 6). The fluctuations in dynamic entropy for novices appear arbitrary both during the resting state and while playing the TOL game, even in low entropy regions.

**Figure 6:**
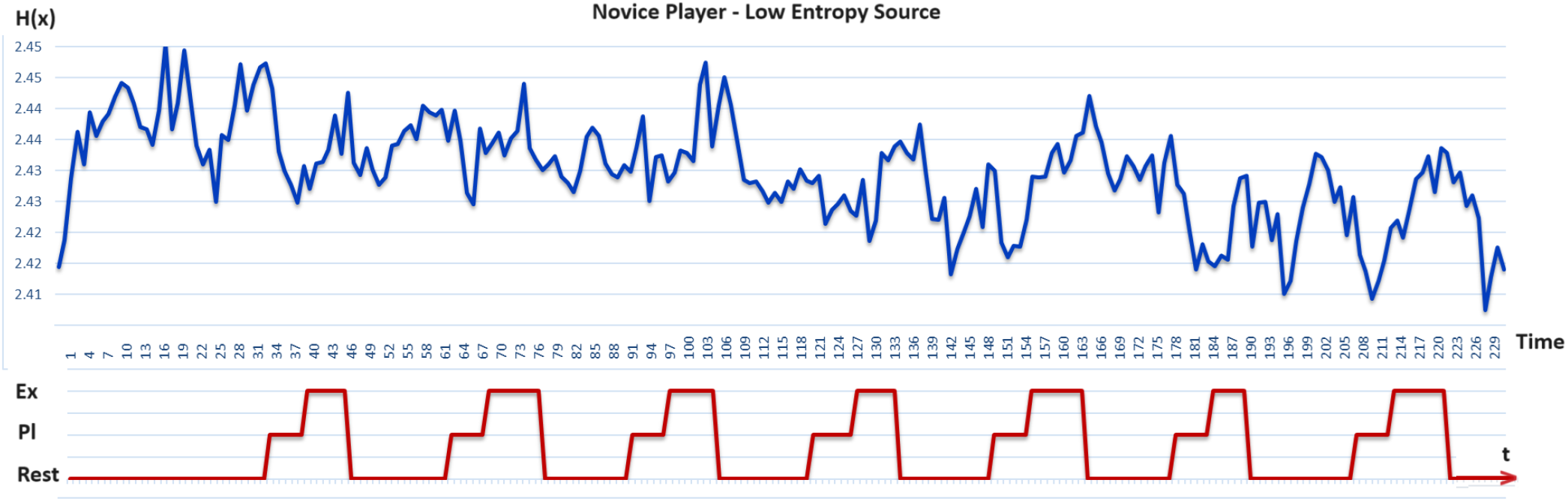
Novice player: Dynamic Entropy fluctuations for a low entropy region, *Right Precuneus* (blue). *Pl* shows the Planning, *Ex* shows the execution and *Rs* shows the resting state time instants (red).

The above observations reveal that dynamic entropy variations during the resting state and game play in relatively low entropy regions provide us with important clues about a player’s degree of expertise.

### 7.3 Low Entropy Information Sources in Complex Problem Solving

The major assumption of this study is that the information sources with the lower entropy are more closely involved in CPS processes compared to higher entropy sources.

During the experiments, we observed that both the static and dynamic entropies are relatively low in the following brain areas:

1. **Frontal lobe**: Superior frontal dorsolateral, Superior frontal orbital, Superior frontal medial, Middle frontal, Precentral gyrus, and Paracentral lobule.
2. **Parietal lobe**: superior and inferior parietal, precuneus, angular gyrus, and post-central gyrus.
3. **Occupital lobe**: Middle occipital, Superior occupital, Inferior occupital, cuneus, calcarine, and lingual gyrus.
4. **Temporal lobe**: Inferior temporal gyrus, Middle temporal gyrus, Fusiform gyrus.

The prefrontal cortex is suggested to be an important part of the cortical network involved especially in planning. (Newman et al., 2003) proposed that left and right prefrontal cortices are equally involved during the solution of moderate and difficult TOL problems. They suggested that the right prefrontal cortex might be connected to the plan generation, while the left part might be more related to the plan execution. The involvement of superior parietal areas is attributed to spatial processing, with the right superior parietal region associated with visuospatial attention necessary for planning. Conversely, the left parietal cortex is linked to visuospatial working memory processing.

In another study, (Newman et al., 2011) suggested that the precentral/inferior frontal junction is implicated in monitoring simple rules and sequence processing. The prefrontal region is proposed to track the retrieval of task-relevant information.

(Lazeron et al., 2000), observed considerable activation in the middle frontal gyrus and the adjacent part of the inferior frontal gyrus, the precentral cortex, and the anterior cingulate gyrus. They revealed that during the TOL planning task, the parietal and occipital regions, including the precuneus, cuneus, and angular gyrus, are actively involved.

Another study, (Fincham Anderson et al., 2002) formalized planning behavior in the Tower of Hanoi (TOH) task as a computational model within the adaptive control of thought–rational (ACT-R) cognitive modeling framework (TOH can be seen as a similar problem-solving measure to TOL). They found that the regions, which differentially responsive to goal-processing operations distribute among the prefrontal cortex, parietal cortex (including the superior parietal, precuneus, cuneus, inferior parietal and angular gyrus), cingulate gyrus, and subcortical structures.

(Unterrainer et al., 2004) proposed that during the TOL task, good problem solvers showed increased activation in the right superior temporal and inferior parietal regions, which may reflect more visuospatial attentional processing. They also observed activation of the right parahippocampal and lingual gyrus during execution, which they suggested reflects the increased demands on spatial working memory. Furthermore, during the execution phase, incorrectly solved trials elicited more widespread activations than correctly solved problems in the premotor and dorsolateral prefrontal cortex, as well as in the left precuneus and the temporal-parietal-occipital regions.

A comparison of our entropy-based approach with the results of the aforementioned experimental studies indicates a significant overlap between low-entropy information sources and active anatomical regions observed in experimental neuroscience for TOL problems. Therefore, we can infer that information sources with low entropy correspond to anatomical regions that are active during complex problem solving.

### 7.4 Static Entropy Analysis for Planning and Execution Subtasks

One of our goals is to compare the activation of brain information sources during the planning and execution subtasks. For this purpose, we estimate the static entropy of each information source on subject, session, and subtask basis. We estimate the static entropy of planning and execution in two sets of experiments.

In the first set of experiments, we use all the voxels to estimate the representative time series for each information source during the planning and execution phases, separately, and, then, estimate the static entropy measures. As it is observed from Figure 7.a, static entropy measures for the planning phase are relatively low compared to that of the execution phase (Gunal Degirmendereli et al., 2019).

**Figure 7:**
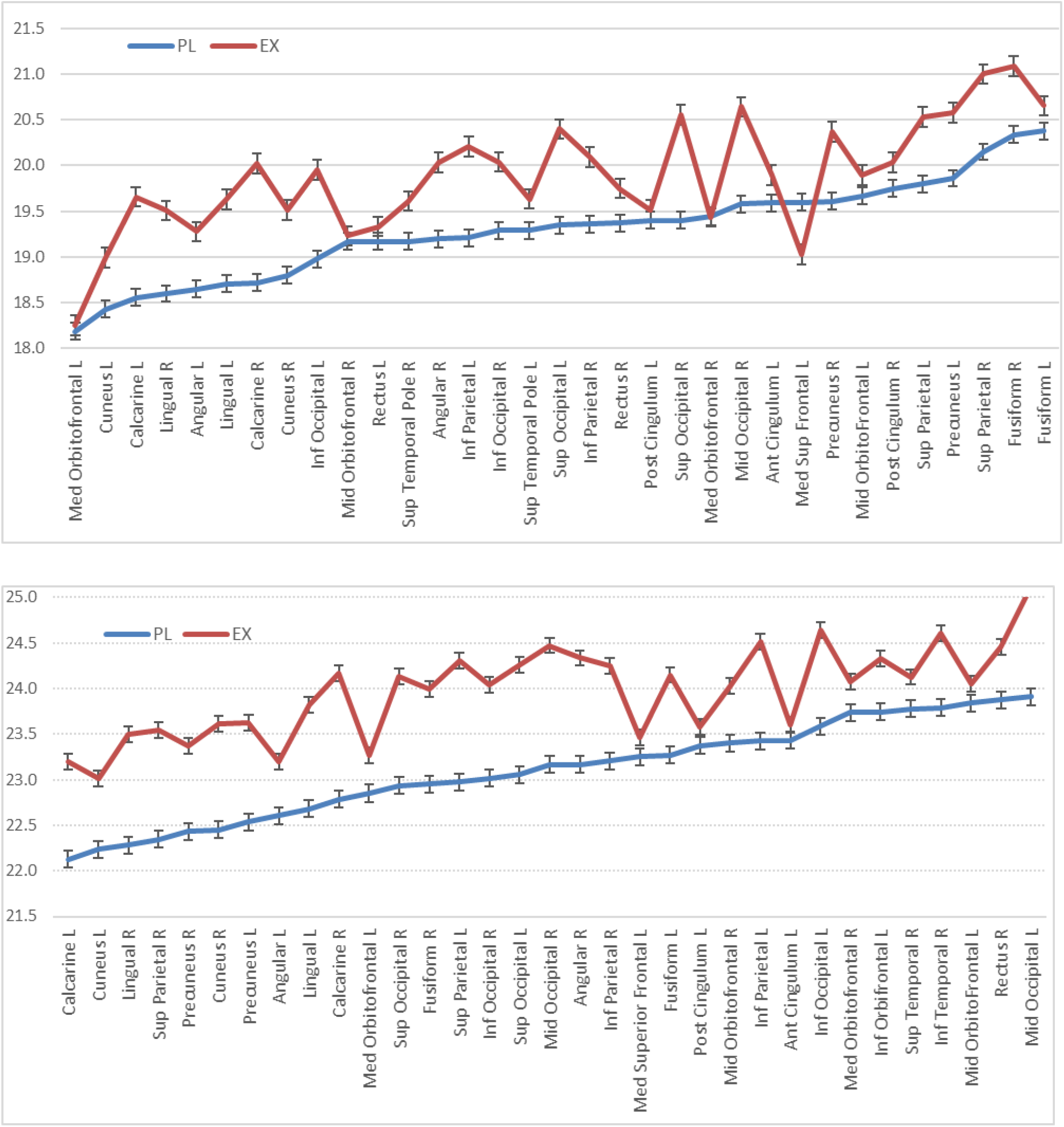
a) The lowest static entropy regions for planning and execution tasks based on all voxels (above). b) The lowest static entropy regions for planning and execution tasks based on the most informative voxels (below).

However, it is well-known that some voxels do not contribute to the underlying cognitive tasks. While these irrelevant voxels may be involved in other brain processes, they can obscure the tasks under investigation by acting as background noise. Various techniques exist in the fMRI literature to eliminate redundant voxels. One popular method is ANOVA, which estimates and ranks the mutual information between a specific cognitive task and the voxel time series. Then, the most informative voxels, with the highest mutual information values are selected from the ranked list and the rest are eliminated. Studies have shown that selecting the most informative voxels using ANOVA significantly improves brain decoding performance for both planning and execution phases (Alchihabi et al., 2018). This finding suggests that removing voxels with low mutual information scores reduces the noise from irrelevant voxels.

In the second set of experiments, we estimate the static entropy values based on the most informative voxels, selected by ANOVA. For this purpose, we estimate mutual information of each voxel time series, when the subject is performing both planning and execution subtasks. Then, we rank the voxels according to their mutual information scores. For each time instant, we select 25.000 voxels out of 185.000. We calculate the representative voxel time series of each information source by averaging only the most informative voxels. Then, we estimate static entropy measures of the planning and the execution phases for all subjects (Figure 7.b)

When we compare Figure 7.a and Figure 7.b, we observe that the difference between the static entropy measures of planning and execution phases is more accentuated when we eliminate the irrelevant voxels. This observation reveals that using only the most informative voxels reduces the static entropy during the planning phase compared to the execution phase. This reduction is due to the elimination of noise from irrelevant voxels.

In the third set of experiments, we applied the same voxel selection and static entropy estimation method to both planning and execution phases, but this time only considering the successful runs. As shown in Figure 8, the difference in static entropy between the planning and execution subtasks is even more pronounced in the successful runs. The planning subtask has much lower entropy measures compared to the execution subtask in all of the low-entropy information sources. The lowest static entropy information sources for planning and execution tasks based on the most informative voxels are visualized in Figure 9

**Figure 8:**
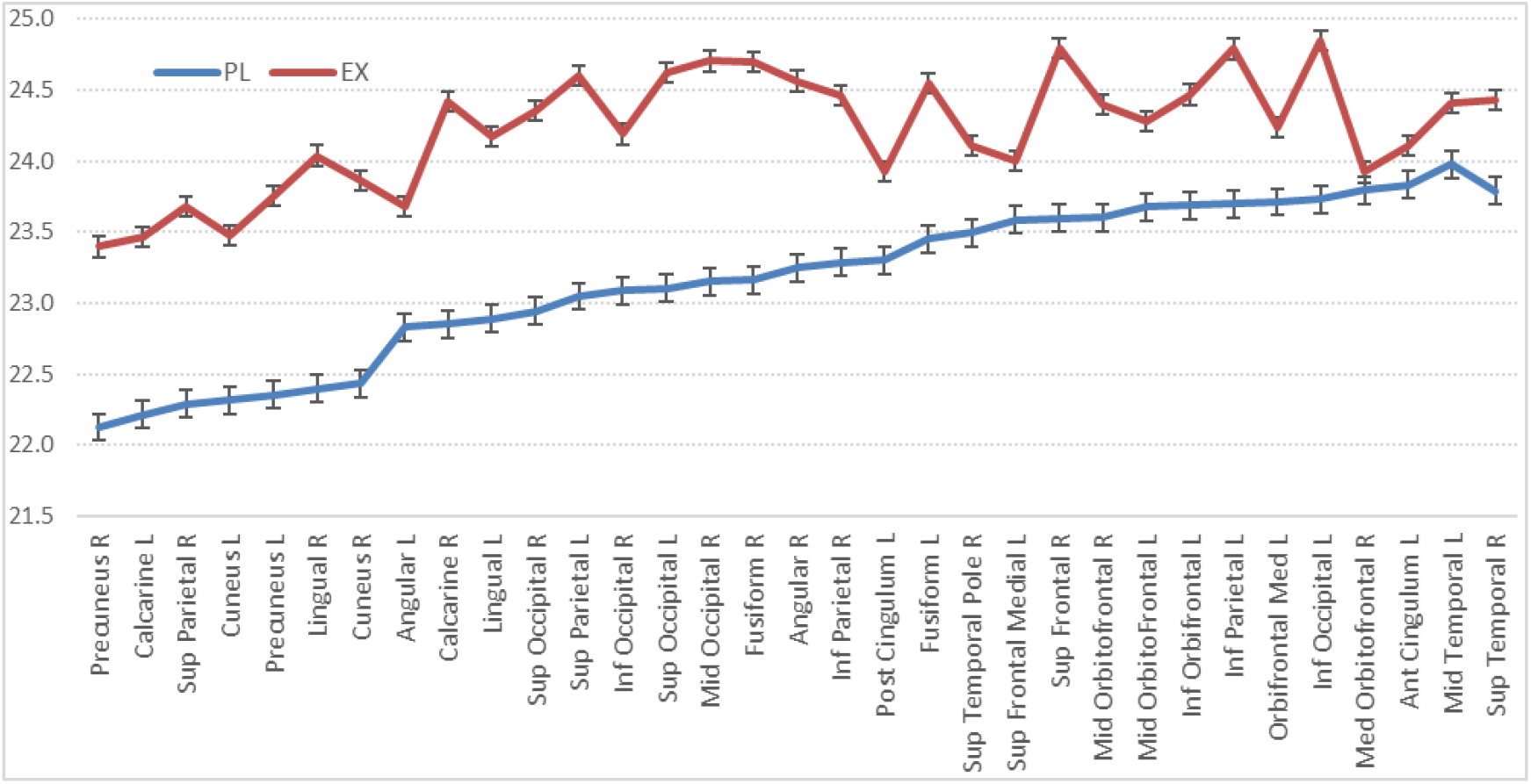
The lowest static entropy regions for planning and execution tasks for successful sessions based on most informative voxels.

**Figure 9:**
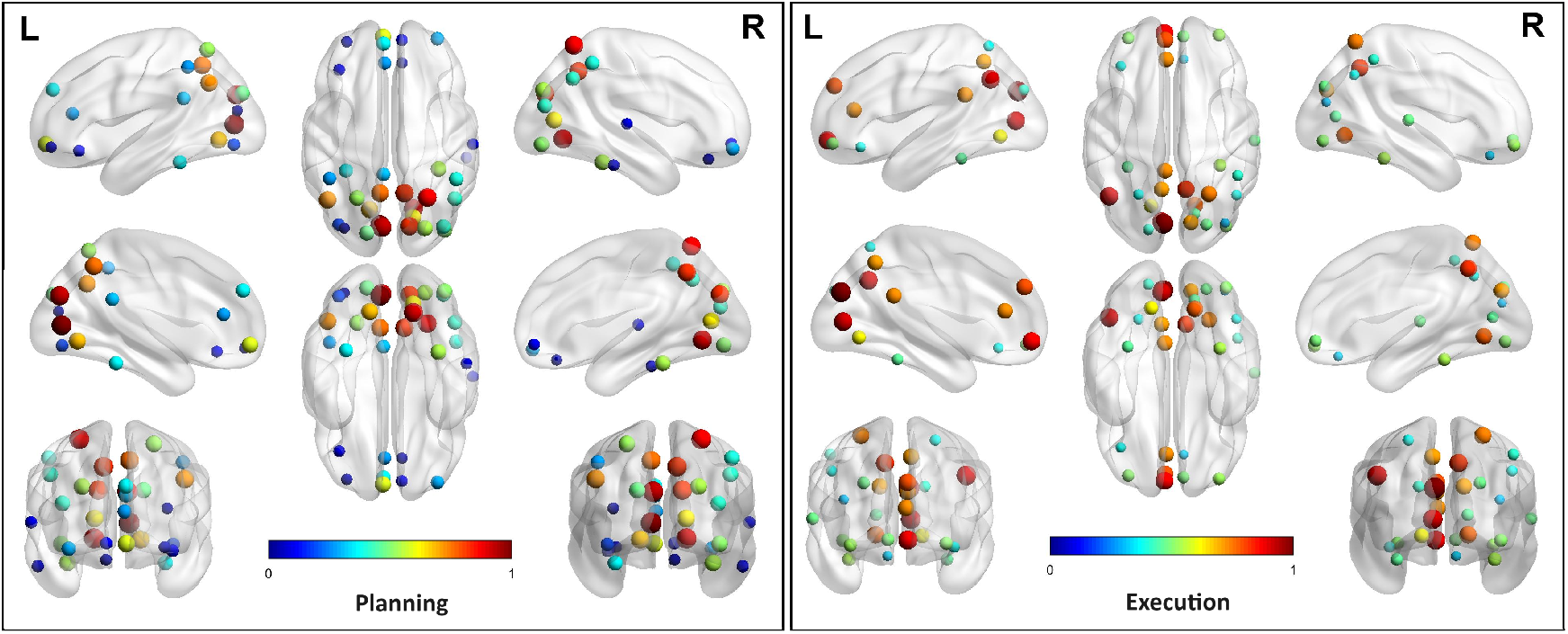
Visualization of anatomical regions, as information sources with the lowest static entropy measures for planning and execution tasks based on most informative voxels. Hot colors represent relatively low static entropy sources. The sizes of the circles are inversely proportional to the static entropy measures

This analysis reveals that the static entropy of the planning task is lower than that of the execution task. The difference between planning and execution entropy is higher for most informative voxels than all voxels. This result is compatible with the previous BOLD analysis study (Alchihabi et al., 2018). Since plan generation requires the participation of numerous cognitive processes, it would lead to more coherent neural processing than the execution task. The difference between planning and execution entropy is higher for successful runs than that for unsuccessful runs. This result can be interpreted as follows: The informative sources of an expert player are more organized compared to that of a novice player.

### 7.5 Static Entropy Analysis for Expert and Novice Subjects

It has been shown that expert and novice players think and solve problems in different ways (Newman & Green, 2015). Experts differ in problem solving skills from novices depending on several factors such as domain-specific knowledge, the quality of memory representations, better short and long-term memory, exploring a broader problem space, elaboration of mental schemas, problem categorization, solution Strategy development, information chunking, and time allocation.

During the Tower of London (TOL) game, a successful planning phase means that a solution plan is effectively developed and stored in working memory, allowing the problem solver to directly implement it. In such cases, the execution task is completed in a relatively short time. However, for complex problems that require multiple subgoals, the solution process often involves an intermix of planning and execution tasks. Consequently, re-planning may be necessary during execution to correct the initial plan, resulting in longer times for the execution-replanning-execution cycle.

In our experiment, we observed that for successful runs, the planning duration was longer than the execution duration. This finding suggests that expert players spent more time planning the solution than implementing it. Conversely, novice players spent less time on planning and more time on executing the plan.

Figure 10 provides a comparison of the low entropy information sources between expert and novice players based on the most informative voxels, selected by ANOVA.

**Figure 10:**
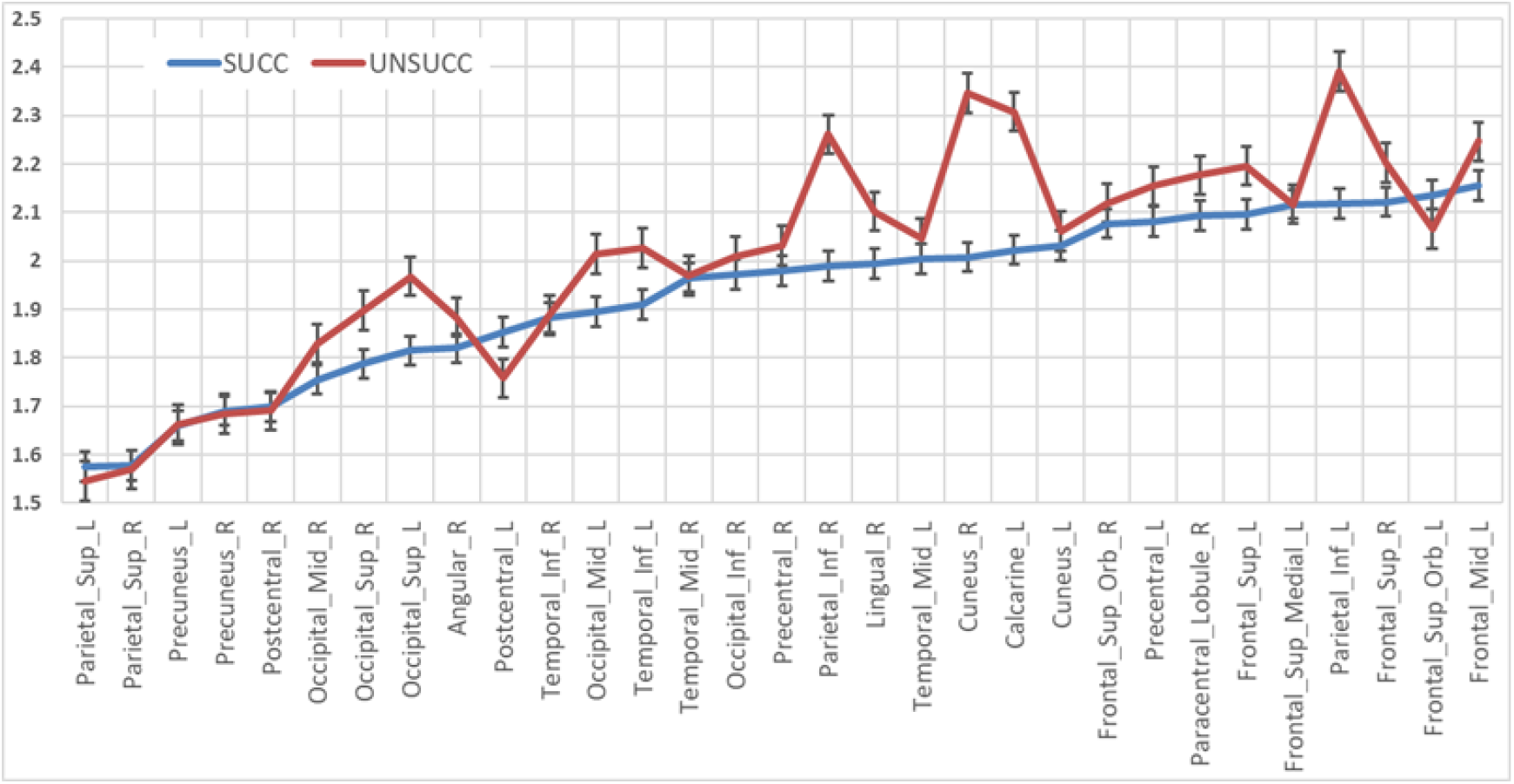
The lowest static entropy regions for successful and unsuccessful sessions based on most informative voxels.

## 8 Experiments on Static Brain Network Representations

In this series of experiments, we estimated the static brain networks introduced in Section 5. Initially, we compared the edge weight matrix of the static brain network with the functional connectivity matrix derived from Pearson correlation. Due to the varying number of voxels in each region, Pearson correlation matrices were calculated based on voxel distributions rather than voxel intensity values.

Our analysis revealed that both matrices exhibited active connections between similar anatomical regions, indicating consistency between the functional connectivity matrices obtained from Pearson correlation and those derived from static KL divergence.

In subsequent experiments, we excluded KL values greater than 0.023 to highlight significant connections and create visually meaningful networks. The static KL matrices were generated separately for the planning and execution phases. In most tasks, we observed that the number of significant connections between information sources was higher during the planning phase than during the execution phase. This finding aligns with established neuroscience research, suggesting that planning is more complex process than execution. (Newman & Green, 2015).

Dynamic brain networks are estimated to examine the behavior of planning and execution tasks in successful and unsuccessful problem solving sessions. We obtained a probability distribution function for each time instant, over all the voxels in an anatomical region and calculated the KL divergences between the regional pairs. Then, we produced separate KL matrices for the planning and the executive time instants of successful and unsuccessful problem solving sessions, and generated functional connectivity graphs.

Figure 11 shows the visualization of information sources that have KL divergence distances less then 0.023 for planning and execution phases for a successful TOL problem solving sessions.

**Figure 11:**
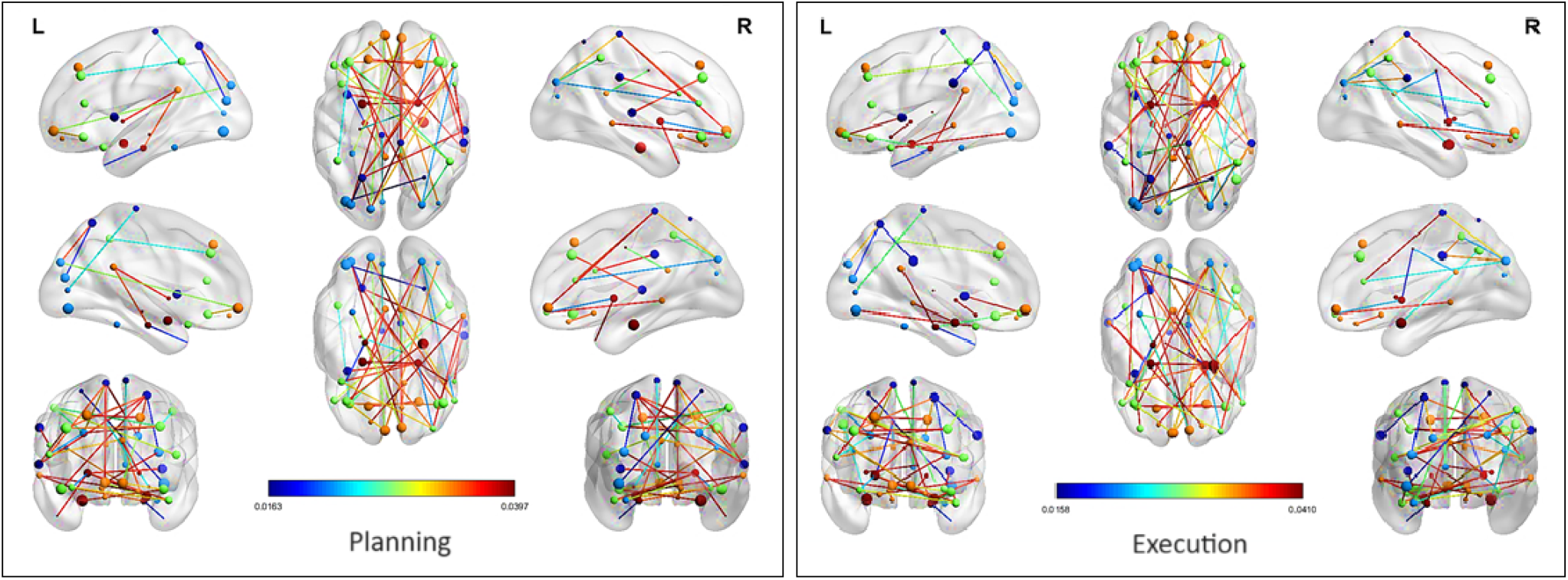
Visualization of the connectivity among anatomical regions, which have shortest KL distance measures for planning and execution tasks for *successful sessions*. The size of the nodes is set according to the node degree (Gunal Degirmendereli, & Yarman Vural, 2021).

Previous studies show that the planning and execution phases of problem solving involve different types of brain activities (Newman & Green, 2015). Planning includes a set of operations for the construction of a problem representation and searching for the appropriate operators to solve the problem. This requires various parts of the neural system to work together. The execution phase involves the implementation of the planned solution, therefore, it requires different neural activities compared to the planning phase. In order to analyze the degree of connectivity among the anatomical regions during the planning and execution phases, we select the most significant KL connections in the edge weight matrices.

Figure 12 shows the visualization of the brain networks that have the smallest KL divergence distances for planning and execution tasks for unsuccessful TOL problem solving sessions. Visual comparison of Figures 11 and 12 reveals the differences in the brain networks of expert and novice players for the planning and execution phases

**Figure 12:**
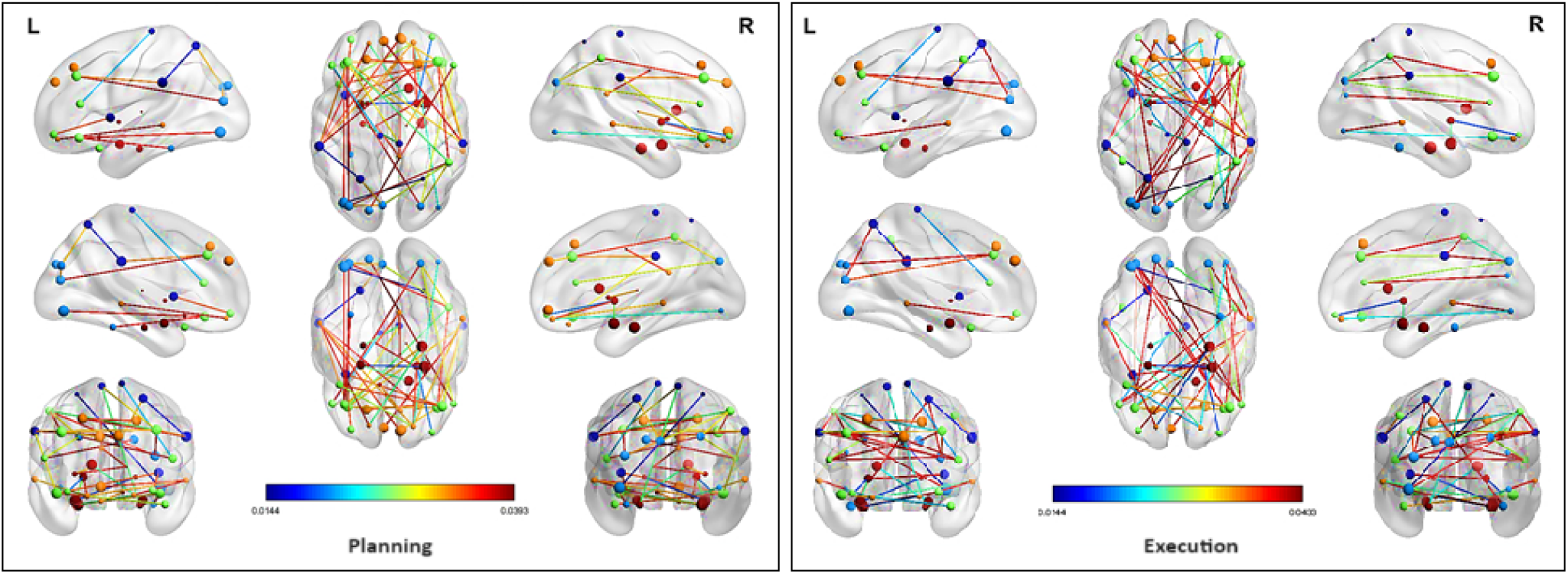
Visualization of the connectivity among the anatomical regions, which have the shortest KL distance measures for planning and execution tasks for *unsuccessful sessions*. The size of the nodes is set according to the node degree (Gunal Degirmendereli, & Yarman Vural, 2021).

## 9 Experiments on Dynamic Entropy and Dynamic Brain Network Representations for Brain Decoding Performances

Two sets of experiments were conducted to evaluate the efficacy of the proposed dynamic entropy and dynamic brain networks with the help of an MLP classifier. The first set evaluates the performance of the MLP in distinguishing the *planning* and *execution* phases, whereas the second set assesses the performance in determining *unsuccessful* and *successful* attempts. The MLP classifier was selected because of its simplicity, efficiency, and ability to comprehend complex brain network patterns.

Two methods were identified for the establishment of a baseline. The first method uses the BOLD signal data as an input to the classifier. Recall that each time instant of the voxel BOLD values of TOL game has either planning or execution phases. We form a feature vector by concatenating the average BOLD signal of each anatomical region at each time instant. Because there are 90 anatomical regions, the input feature vectors have a dimension of 1 *×* 90, each of which has a label of planning or execution. In the second baseline, we estimate the Pearson correlation coefficients between the representative time series of anatomical regions. Thus, the input space consists of 90 *×* 90 = 8100 dimensional feature vectors

We tested the dynamic entropy and dynamic brain network representations compared to the baselines. The entropy data consists of feature vectors of 90 values, each corresponding to an estimation of the first-order dynamic entropy of an anatomical region. On the other hand, the KL data consists of 60 *×* 60 matrices, where 30 high-entropy regions were eliminated to reduce noise from irrelevant regions and avoid the curse of the dimensionality problem.

All the experiments were performed using 10-fold cross-validation, with 10% of the dataset as a validation set for each fold. Additionally, we employ a wide range of performance metrics, including loss, F1 score, precision, and accuracy, to quantify the results. With the help of these approaches, we ensure the reliability of the results and allow for comprehensive performance comparisons.

In the planning-execution classification experiment, the KL feature vectors achieved the highest performance across the metrics with an accuracy of 0.9104 ± 0.0062. Following this, the Pearson correlation vectors attained an accuracy of 0.9045 ± 0.0066 and an F1 score of 0.9044 ± 0.0066. The entropy and BOLD signal vectors showcased the second-lowest and lowest performances with accuracy scores of 0.8589 ± 0.0189 and 0.7506 ± 0.1065, respectively.

In the successful-unsuccessful classification experiment, the KL feature vectors again achieved the highest performance with an accuracy of 0.9854 ± 0.0071, loss of 0.0336 ± 0.0150, and F1 score of 0.9852 ± 0.0072. Like the previous experiment, Pearson, entropy, and BOLD features ranked second, third, and fourth across all the metrics.

It can be seen that the KL feature vectors consistently performed the best in both experiments. The KL and entropy vectors performed better than the Pearson and BOLD vectors, respectively, demonstrating their ability to capture complex brain activities during TOL tasks. The overall performance of the KL and Pearson vectors is better than that of the entropy and BOLD vectors. This gap can be explained by the larger vector size of 90 *×* 90 compared to only 90 elements.

**Table 1:** Classification performances of MLP for planning and execution tasks.

| Data | Loss | Accuracy | Precision | Recall | F1 |
| --- | --- | --- | --- | --- | --- |
| KL | **0.2163 ± 0.0133** | **0.9104 ± 0.0062** | **0.9107 ± 0.0065** | **0.9105 ± 0.0062** | **0.9103 ± 0.0063** |
| Pearson | 0.2218 ± 0.0157 | 0.9045 ± 0.0066 | 0.9054 ± 0.0061 | 0.9050 ± 0.0064 | 0.9044 ± 0.0066 |
| Entropy | 0.3343 ± 0.0334 | 0.8589 ± 0.0189 | 0.8663 ± 0.0145 | 0.8577 ± 0.0209 | 0.8575 ± 0.0204 |
| BOLD | 0.6811 ± 0.5008 | 0.7506 ± 0.1065 | 0.7733 ± 0.1739 | 0.7468 ± 0.1129 | 0.7174 ± 0.1579 |

**Table 2:** Classification performances of MLP for successful and unsuccessful attempts.

| Data | Loss | Accuracy | Precision | Recall | F1 |
| --- | --- | --- | --- | --- | --- |
| KL | **0.0336 ± 0.0150** | **0.9854 ± 0.0071** | **0.9851 ± 0.0073** | **0.9853 ± 0.0071** | **0.9852 ± 0.0072** |
| Pearson | 0.0543 ± 0.0203 | 0.9752 ± 0.0076 | 0.9752 ± 0.0069 | 0.9748 ± 0.0083 | 0.9749 ± 0.0077 |
| Entropy | 0.0788 ± 0.0238 | 0.9652 ± 0.0082 | 0.9649 ± 0.0083 | 0.9656 ± 0.0074 | 0.9649 ± 0.0082 |
| BOLD | 0.1836 ± 0.0776 | 0.9142 ± 0.0338 | 0.9209 ± 0.0265 | 0.9110 ± 0.0365 | 0.9122 ± 0.0358 |

## 10 Discussion / Limitations

### 10.1 Entropy Estimation and Sample Size Considerations

One methodological consideration concerns potential bias in entropy estimation due to limited sample sizes when estimating PDFs from voxel intensities within anatomical regions. Nonparametric entropy estimation methods such as KDE are known to exhibit bias when sample sizes are small. However, our dataset includes anatomical regions with voxel counts ranging from 220 to 5104, which provides a statistically sufficient sample size for stable estimation of a univariate density estimation.

Theoretical work (Paninski, 2003) demonstrates that entropy estimation bias decreases inversely with sample size (O(1/N)). To further control estimation bias, we employed region-specific adaptive bandwidths (Silverman, 1986). Although fMRI voxel intensities exhibit spatial autocorrelation, previous research (e.g., Wang et al., 2014) supports the application of KDE to voxelwise entropy estimation even in the presence of spatial dependencies.

To empirically assess the potential impact of voxel count, we examined the relationship between average static entropy and region size. As shown in Figure 13, there is no monotonic relationship between voxel count and entropy, suggesting that sample size does not trivially drive entropy estimates.

**Figure 13:**
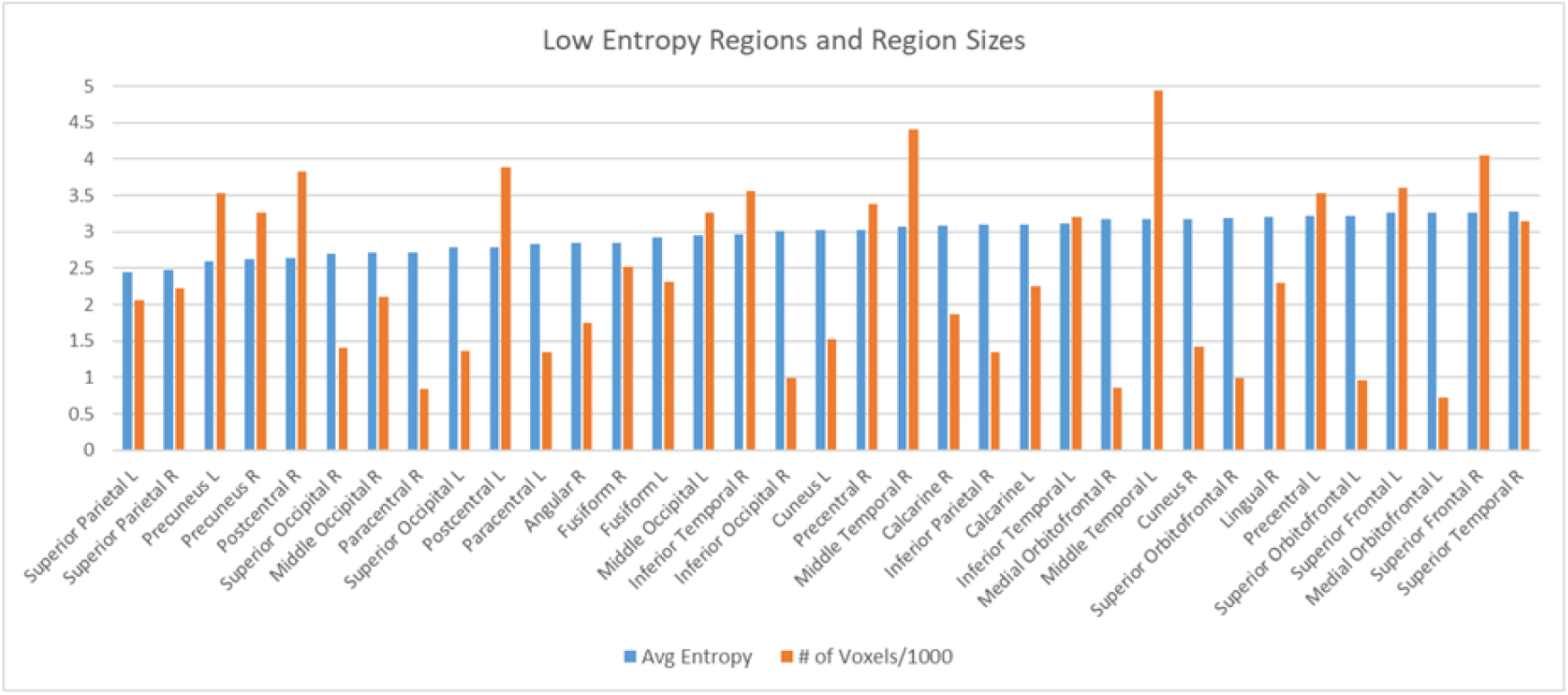
The relationship between average regional static entropy and region size for low-entropy regions.

While the present study focused on spatial entropy derived from voxel distributions at individual time points, future studies will incorporate time-resolved entropy measures, such as sliding-window entropy and multiscale entropy, to capture temporal complexity within regions. Such extensions would complement the current spatial framework by capturing how internal signal regularity evolves over time. We also plan to explore bias-corrected entropy estimators (e.g., Nemenman–Shafee–Bialek (NSB) entropy) to further enhance robustness.

### 10.2 Statistical Validation of KL Divergence

One limitation of the current study is the absence of formal permutation-based statistical validation for KL divergence values. Ideally, significance would be assessed by generating null distributions through shuffling or permutation of regional time series. However, the computational demands of our dataset (72 sessions × 90 regions × 590 time points) render comprehensive permutation testing infeasible at present.

To partially address this limitation, we compared KL divergence-derived connectivity matrices to Pearson correlation matrices computed from the same voxel distributions. Despite differences in the nature of these metrics, we observed notable structural overlap among the strongest connections, suggesting that KL Divergence networks reflect meaningful functional organization (Figure 14).

**Figure 14:**
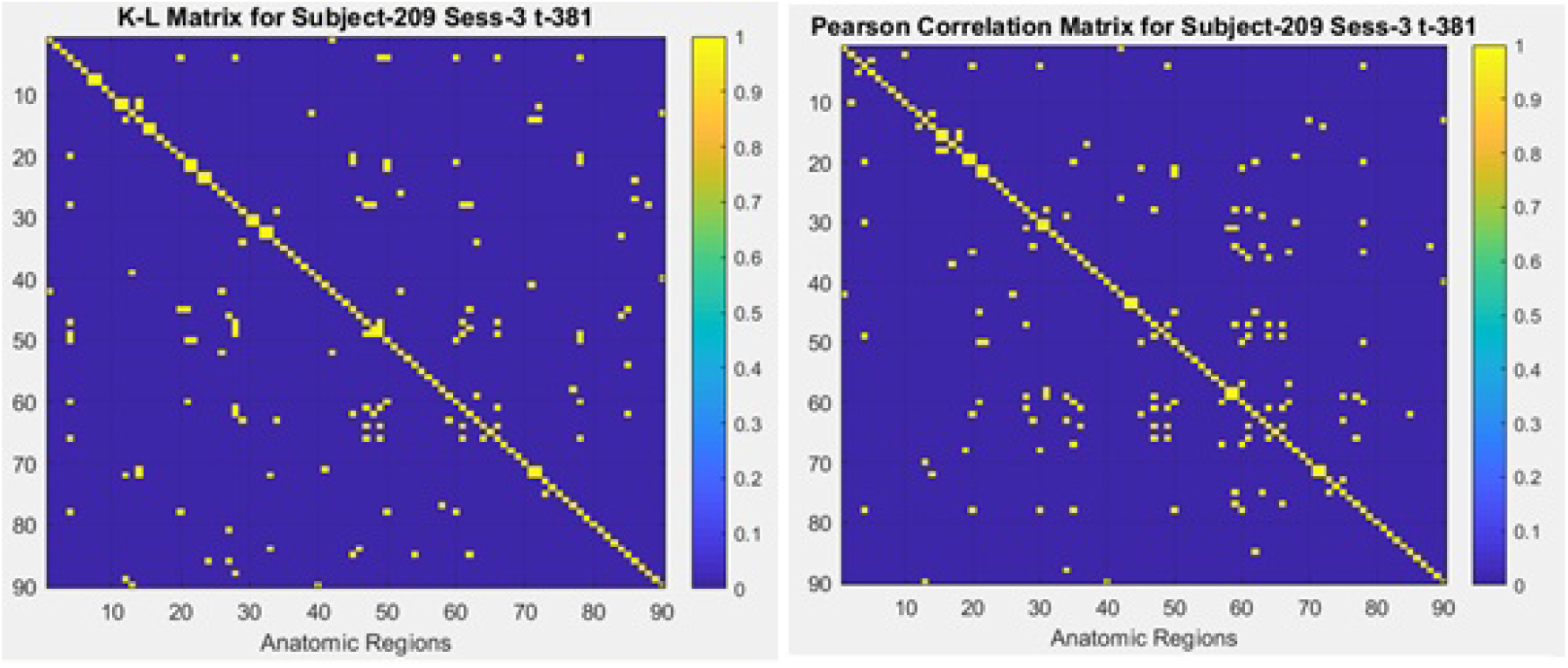
Comparison of thresholded KL Divergence (left) and Pearson correlation matrices (right) at a representative time point (Subject 209, Session 3, Time Instant 381).

Nonetheless, we acknowledge that direct null model validation is necessary for full statistical rigor. As a future work, the statistical stability of the suggested methods will be further analyzed by null modeling, which will strengthen the statistical interpretation of dynamic information-theoretic connectivity networks derived from fMRI.

## 11 Conclusion

This study presents an enhanced information-theoretic framework for analyzing functional brain activity using entropy-based measures and Kullback-Leibler (KL) divergence network models. We propose a computational model based on Shannon information theory to estimate active brain regions and their interactions during complex problem-solving tasks.

To characterize information processing in the brain, we model each anatomical region as an information source, where dynamic entropy captures temporal fluctuations in neural activity, and static entropy quantifies the overall information content within each region. This dual-entropy framework enables the investigation of both transient and stable informational properties of brain function.

Applying our model to fMRI data from Complex Problem Solving (CPS) tasks, we analyze the distinct contributions of brain regions across different cognitive states. We construct static and dynamic brain networks to represent inter-regional interactions during the main phases of CPS. Our model also provides a basis for comparing expert and novice players by analyzing entropy variations during the planning and execution phases of the CPS task.

A comparison with experimental studies in the literature reveals a significant overlap between low-entropy anatomical regions and those identified as active in neuroscience research on the Tower of London (TOL) task. These results support the use of entropy-based metrics to identify functionally engaged brain areas. Regions with lower entropy exhibit increased neural activity, consistent with prior research. Furthermore, KL divergence-based network models provide a robust framework for capturing dynamic brain interactions, achieving over 90% classification accuracy in decoding cognitive states.

Antonacci et al. (2025) emphasized that nonstationary neural processes require dynamic information-theoretic measures, rather than static correlations, to decode rapid cognitive transitions. Our findings align with this perspective: we demonstrate that time-resolved KL Divergence networks derived from fMRI can effectively distinguish between planning and execution phases of problem solving—even within the temporal constraints of fMRI. By combining the anatomical precision of fMRI with a time-sensitive information-theoretic framework, our approach bridges a critical gap in mapping cognitive flexibility across large-scale brain networks.

Our findings offer new insights into how brain regions coordinate during complex cognitive tasks and introduce a novel computational tool for analyzing dynamic brain network activity. The proposed framework may contribute to future research in cognitive neuroscience, neuroinformatics, and brain-computer interface development.

Information theory serves as a powerful bridge between the mind and brain, enabling a deeper understanding of cognition. By revealing the neural basis of specific cognitive tasks, our study contributes to a more precise characterization of high-level cognitive processes, such as complex problem solving.

## Acknowledgment

We thank Professor Sharlene Newman for providing us the TOL data. The work is supported by TUBITAK (Scientific and Technological Research Council of Turkey) under the grant No:116E091.

